# Halogen bonds between ligands and proteins: can we use them in validation?

**DOI:** 10.1101/2025.06.24.661284

**Authors:** Ida de Vries, Georgia Tsiompanaki, Anastassis Perrakis, Robbie P. Joosten

## Abstract

Halogen bonds are polar interactions between a halogen atom and its acceptor. They have characteristic geometry induced by the positively charged σ-hole on the halogen atom. Despite their importance in drug development, halogen bonds are often overlooked in the analysis and validation of ligand-protein complexes.

We analysed halogen bonds between ligands and proteins in structure models from the PDB-REDO databank and defined key geometric parameters. These are the donor-acceptor distance, and two bond angles (θ1 and θ2) for interatomic halogen bonds, or in the case of halogen-π interactions, the distance between the halogen and acceptor π-system and a bond angle (θ1).

Based on the distribution of these geometric parameters, we introduce a score, HalBS, that marks whether halogen bonds are adopting the preferred or allowed geometry or are an outlier and should be examined critically. A reference implementation of this score is now available in PDB-REDO. This is a first step towards improving halogen bond treatment in the analysis of macromolecular structure models.

## Introduction

Halogen bonds are polar interactions between a halogen atom and an acceptor atom. A halogen bond occurs due to polarisation of the σ-bond between the halogen and its connecting atom. This causes a positive charge on the opposite site of the halogen atom, the so-called σ-hole (Figure 1A) (Clark et al. 2007). As a result, the acceptor of the halogen bond is typically an atom with a free lone-pair, like an oxygen, nitrogen, sulfur, phosphorus or selenium atom. Additionally, halogen atoms can interact with π-systems (Bent 1968). Due to differences in polarisation properties, halogen bonds with iodine are the strongest, followed by bromine, chlorine and finally halogen bonds with a fluorine atom are the weakest halogen bonds (Politzer et al. 2010; Gilday et al. 2015; Cavallo et al. 2016).

**Figure 1.**
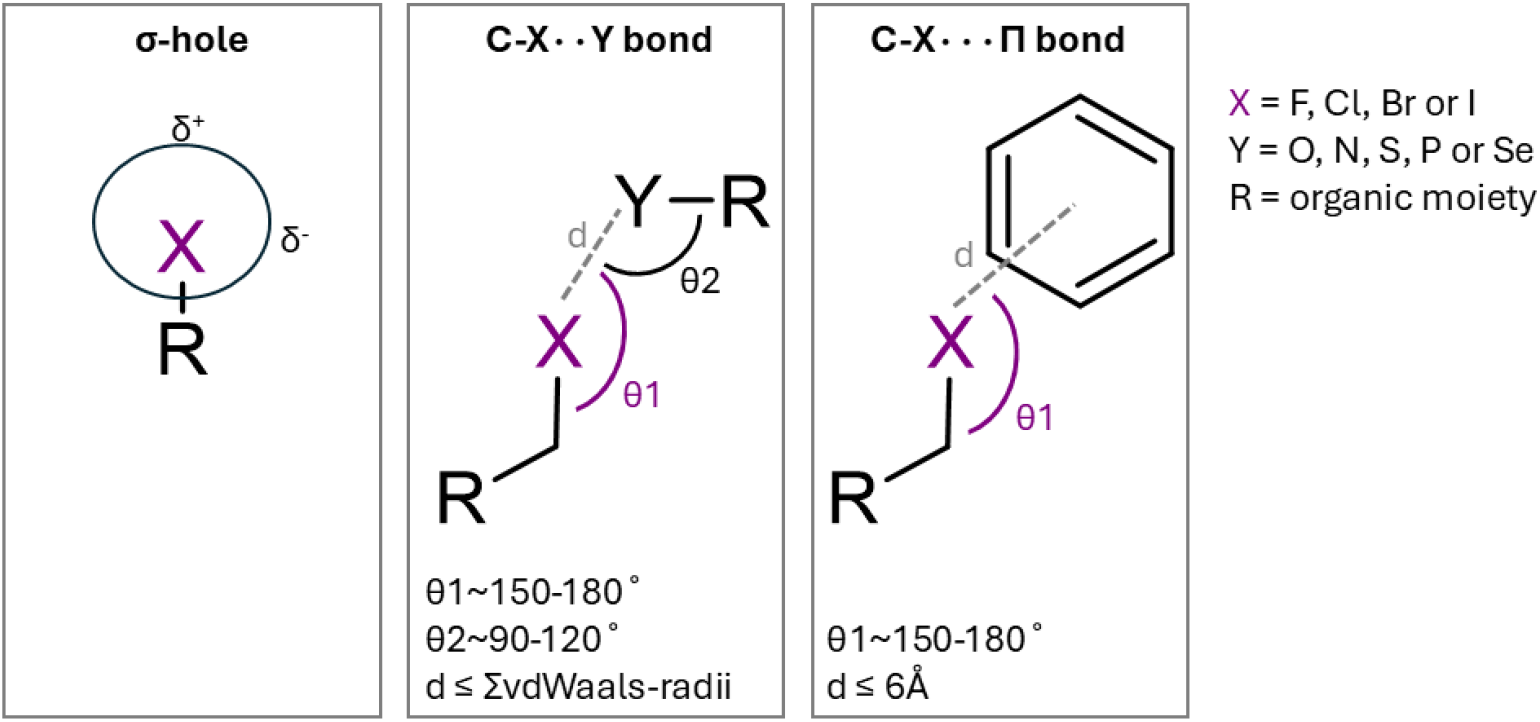
Schematic representation of the σ-hole on a halogen atom (left), and halogen bond characteristics of C-X··Y halogen bonds (middle) and C-X⋯Π halogen bonds (right).

Halogen bonds can be characterized based on key geometric parameters. For the “regular” interatomic halogen bond, in this work referred to as C-X··Y halogen bonds, the first parameter is the distance (d) between the halogen and acceptor atom. As halogen bonds are attractive interactions, the distance can be shorter than the sum of the Van der Waals radii of the atoms (Hassel 1970; Auffinger et al. 2004). For the orientation of the acceptor with respect to the halogen atom two angles are defined: θ1, the angle between the halogen containing compound and the acceptor, and θ2, the angle between the halogen atom and the acceptor containing compound (Figure 1B). These angles show distinct ranges, caused by the position of the σ-hole of the halogen atom. For C-X··Y halogen bonds ideally θ1 spans between ∼150° and ∼180° and θ2 between ∼90° and ∼120° (Bent 1968; Desiraju et al. 2013; Scholfield et al. 2013; Cavallo et al. 2016; Turunen et al. 2021). In the case of halogen bonds between a halogen atom and a π-system, the halogen-π interaction, in this work referred to as C-X*⋯*Π halogen bonds (Figure 1C), only the distance between the halogen atom and the π-system, and the θ1 angle are defined (Desiraju et al. 2013; Cavallo et al. 2016; Jubb et al. 2017; Kellett et al. 2020).

Halogen atoms and their potential to form halogen bonds are used in drug design, as halogens affect the affinity, selectivity and efficacy of therapeutics (Kortagere et al. 2008; Lu et al. 2009; Lu et al. 2010; Fraley and Sherman 2018). Halogen substituents increase the overall hydrophobicity of a ligand, thereby enhancing the hydrophobic effect of binding, driven by the minimization of unfavourable water–nonpolar contacts upon binding, and yielding a favourable entropic contribution to the binding free energy (Langton et al. 2016). Thus, halogenation of ligands alters the conventional hydrophobic effect by combining the nonspecific entropic gain of water release with the specific directional interactions inherent to halogen bonds (Wang et al. 2019; Verteramo et al. 2024).

These properties make halogens attractive features in drug design but also in the earlier, more explorative drug lead discovery phase. Herein, halogens are often found in molecules used in virtual and fragment screening libraries, used to discover potential new binders to a target protein (Heidrich et al. 2019; Dammann et al. 2022; Chopra et al. 2023). Particularly, libraries that include fluorine containing compounds are especially valuable in fragment screening using Nuclear Magnetic Resonance (NMR) (Troelsen et al. 2020; Li and Kang 2024). Fluorine atoms are not abundant in endogenous molecules but are favourable for NMR due to their high sensitivity and wide chemical shift range (Buchholz and Pomerantz 2021). Hence, fragment screening NMR is typically used to discover protein binding sites and protein dynamics (Danielson and Falke 1996; Li and Kang 2024). They can also be used to identify compounds that bind to RNA (Kreutz et al. 2006).

Despite their overall benefits for drug design and discovery, halogen bonds are often overlooked in macromolecular structure model validation (Zhang et al. 2017). One of the few places where halogen bonds are explicitly considered is at the PDBe-KB (Protein Data Bank in Europe Knowledge Base) (PDBe-KB consortium 2022; Choudhary et al. 2025) pages for ligands, where the determination of halogen bonds originates from Arpeggio (Jubb et al. 2017). Here, we analyse halogen bonds found in PDB-REDO structure models and propose validation metrics for halogen bonds between a halogen containing ligands and amino acid residues. in the protein. Using such metrics, halogen bonds can be incorporated and integrated into pipelines that validate protein-ligand interactions.

## Results and Discussion

### Defining halogen bonds between ligands and proteins in PDB-REDO

To define halogen bonds in macromolecular structure models, we selected every fluorine (F), chlorine (Cl), bromine (Br) and iodine (I) atom present in PDB-REDO structure models. These were stored together with their most likely acceptor atoms (i.e. O, N, S, P or Se). We used the structure models from the PDB-REDO databank to ensure all models are uniformly treated and no specific restraints for halogen bond interactions were used (Joosten et al. 2009). To ensure high quality data, we selected halogen atoms with a B-factor equal to or below 100, a Real Space Correlation Coefficient (RSCC, (Jones et al. 1991)) for the compounds containing the halogen or acceptor atom equal to or above 0.9 which were in structure models with a resolution equal to or better than 2.5Å (complete distributions in supplemental Figure S1).

Halogen bonds can be formed with different acceptor atoms and residues in proteins. Backbone oxygen atoms, but also oxygen atoms in the side chains of amino acids can serve as acceptor atoms. Furthermore, nitrogen atoms with a free lone pair and sulphur atoms in amino acid side chains can fulfil this role (Wilcken et al. 2013). Regarding the nitrogen atom as acceptor, only histidine residues are, depending on the context-specific protonation state, potential acceptors within a protein. However, within our initial data we did not find high-confidence observations, hence nitrogen atoms were excluded for the remainder of this study.

To be able to differentiate between acceptors for C-X··Y bonds in macromolecular structure models, we mined the acceptor with its neighbouring connecting atom. For example, if the acceptor is an oxygen atom of a hydroxyl group on a carbon atom, the carbon atom is stored with the oxygen atom and the acceptor is marked as ‘O-C’. The acceptors of the found halogen bonds mainly appear to be O-C acceptors, followed by S-C. There are a few observations for acceptor atoms attached to a non-carbon atom (supplemental Table S1), which for example occurs when the acceptor is a non-canonical amino acid. These were too few observations to define sensible validation targets for these acceptors. Therefore, only O-C and S-C acceptors were finally selected for further analysis of C-X··Y bonds.

For the halogen bonds involving a π-system, the side chains of histidine, phenylalanine, tyrosine, and tryptophan are acceptor candidates (Wilcken et al. 2013). We mined these to identify C-X*⋯*Π halogen-acceptor pairs in PDB-REDO structure models. Halogen bonding interactions with the π-system of phenylalanine are found most (3481), followed by tyrosine (2442), histidine (1549), and tryptophan (935) (supplemental Table S1). We then selected only the halogen bonds in which the halogen atom is in a ligand, i.e. any compound marked as ‘non-polymer’ in the CCD, (supplemental Table S2) and the acceptor is a canonical amino acid in a protein.

The final dataset contains 8,423 C-X··Y and 8,096 C-X*⋯*Π ligand-protein halogen bonding interactions (Table 1), which were used in downstream analyses.

**Table 1:** Number of observations of halogen-acceptor pairs of ligand-protein interactions for C-X··Y and C-X⋯Π bonds found in PDB-REDO structure models. For C-X··Y bonds the acceptor is listed with its neighbouring atom, e.g. O-C is an oxygen atom as acceptor, connected to a carbon atom. Accepting π-systems are listed by amino acid.

| Number of halogen-acceptor pairs in ligand-protein interactions |  |  |  |  |  |
| --- | --- | --- | --- | --- | --- |
| Halogen<br>Acceptor | F | Cl | Br | I | Total |
| **C-X...Y halogen bonds** |  |  |  |  | **8423** |
| **O-C** | 4381 | 456 | 2479 | 156 | 7472 |
| **S-C** | 554 | 80 | 312 | 5 | 951 |
| **C-X...$\pi$ halogen bonds** |  |  |  |  | **8096** |
| **Phenylalanine** | 2097 | 167 | 1100 | 49 | 3413 |
| **Tyrosine** | 1328 | 102 | 895 | 14 | 2339 |
| **Histidine** | 1135 | 55 | 257 | 9 | 1456 |
| **Tryptophan** | 539 | 121 | 225 | 3 | 888 |

### The geometry of ligand-protein halogen bonds in PDB-REDO

For the ligand-protein halogen bonds in the dataset, we calculated the characteristic halogen bond geometries: distance, Van der Waals overlap, and the θ1 and θ2 angles for C-X··Y halogen bonds; and distance and θ1 angle for C-X*⋯*Π halogen bonds. We then investigated the distribution of these metrics and visualized them as boxplots (Krzywinski and Altman 2014), unless there were 25 or fewer observations, in which case individual observations are shown (Figure 2).

**Figure 2.**
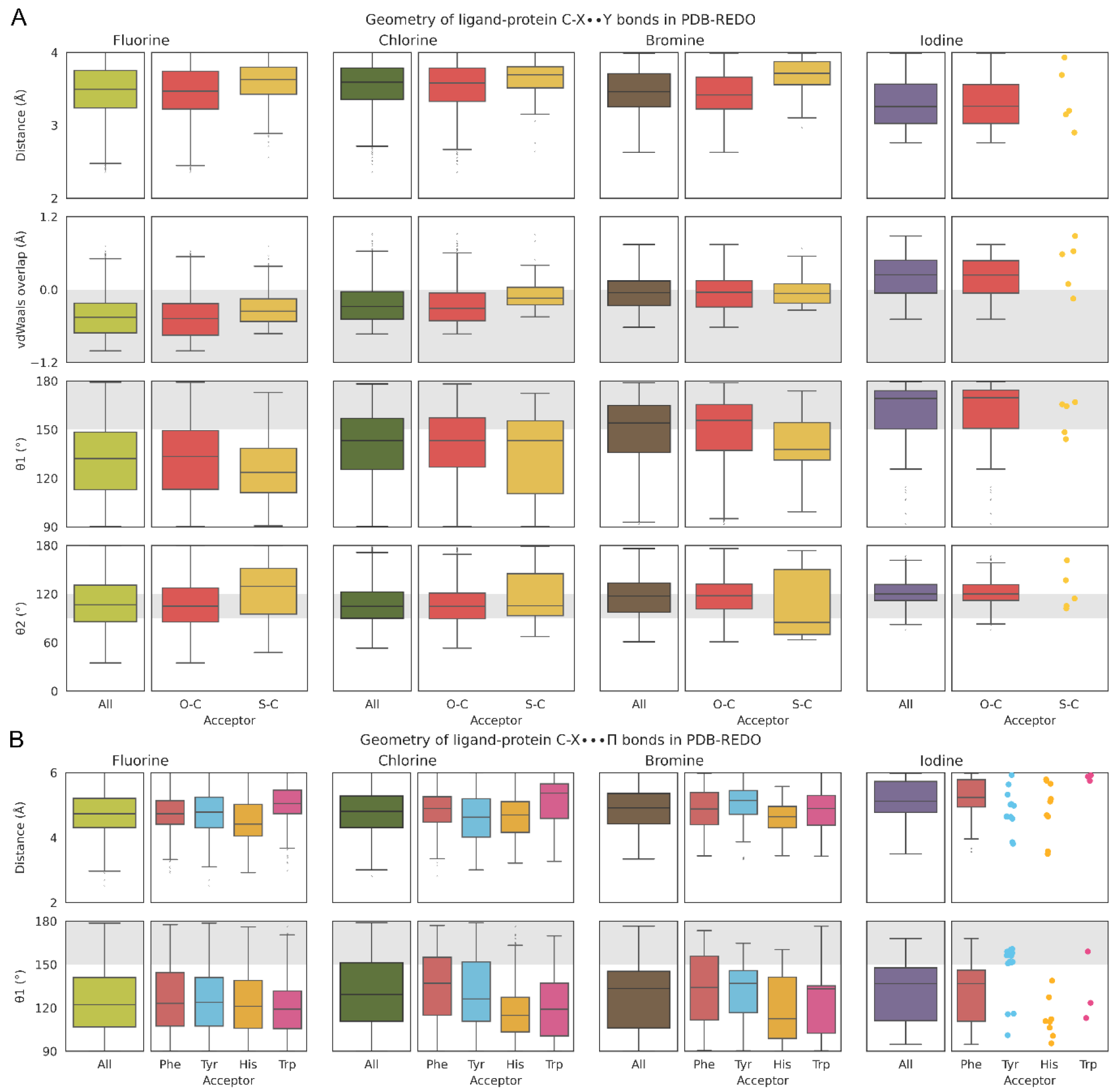
A) Geometry of ligand-protein C-X··Y halogen bonds found in PDB-REDO. For each halogen (from left to right: fluorine (light green), chlorine (dark green), bromine (brown), iodine (purple)) the overall distance, Van der Waals overlap, θ1 and θ2 angles are shown on the left panel. In the panels on the right-hand side these values are split per acceptor: Oxygen (red, O-C), and Sulfur (yellow, S-C). Note that a positive value for the Van der Waals overlap means that the distance between halogen and acceptor atom is shorter than sum of the Van der Waals radii. B) Geometry of ligand-protein C-X⋯Π halogen bonds found in PDB-REDO. For each halogen (from left to right: fluorine (light green), chlorine (dark green), bromine (brown), iodine (purple)) the overall distance and θ1 angle are shown on the left panel. In the panels on the right-hand side these values are split by acceptor: the π-system in the side chains of histidine (yellow), phenylalanine (red), tyrosine (blue) and tryptophan (pink). Grey areas indicate the expected ranges of the geometries. Boxes span from first to third quartile, the median is depicted as the middle line and whiskers extend to 1.5 times the interquartile range (IQR). Statistical values corresponding to the boxplots (number of observations, mean, median, Q1, Q3 and Median absolute deviation (MAD)) can be found in Supplemental Tables S3 and S4 for panels A and B, respectively. For clarity a strip plot is created when there are 25 or fewer observations.

From the boxplots for C-X··Y halogen bonds (Figure 2A), we observe that the distribution of the distance of the halogen bond is rather narrow (around 3.5Å), independent of halogen type or acceptor atom. Note that the median distance becomes shorter and the distribution broader for halogen bonds involving iodine. A Van der Waals overlap between the halogen atom and its acceptor is barely present for fluorine and chlorine. Notably, for bromine and even more so for iodine, halogen-acceptor pairs do show an overlap with their Van der Waals radii. When investigating the θ1 angle for protein-ligand C-X··Y halogen bonds, we observe smaller median angles than the expected range of 150° - 180°. Particularly halogen-acceptor pairs involving fluorine (132°), chlorine (143°) have smaller median θ1 angles, while for bromine (154°) and iodine the median θ1 angle (169°) is within the expected range. Contrary to the θ1 angles, the median θ2 angles are within their expected range of 90° - 120°, independent of the halogen type.

For C-X*⋯*Π halogen bonds (Figure 2B), we observe that the median distance of the halogen bond is rather stable (around 4.8Å) for fluorine, chlorine and bromine, while the halogen bonds involving iodine are longer (median 5.1 Å). The observed θ1 angles for the halogen-π bonds have a median around 110°-140°, whereas we would expect them to range between 150° and 180°. While the distances between halogen and acceptor π-system is not substantially different for different acceptor amino acids, the θ1 angles show different preferences. Particularly the histidine and tryptophan residues seem to have a smaller θ1 angle in halogen bonds with chlorine and bromine.

### Validation of halogen bonds

For the C-X··Y halogen bonds, we observe increasing Van der Waals overlap for halogen-acceptor pairs with increasing halogen atom size (fluorine < chlorine < bromine < iodine). Following a similar trend, the θ1 and θ2 angles reach their expected values with increasing halogen atom size. This fits the increasing polarization of the σ-hole and the electron negativity of halogen atoms (Politzer et al. 2010; Cavallo et al. 2016), resulting in strongest halogen bonds involving iodine and weakest for fluorine atoms. Hence, halogen interaction pairs involving fluorine or chlorine should be interpreted with care and might not be considered ‘true’ halogen bonds. On the other hand, halogen-acceptor pairs with bromine and iodine are more in line with theory and hence can be considered ‘true’ halogen bonds. Note that C-X··Y halogen bonds involving bromine show little Van der Waals overlap, but the θ1 and θ2 angles are within the expected ranges. Halogen bonds involving iodine are even more in line with “textbook” halogen bonds: they show Van der Waals overlap, their θ1 angle nicely ranges between 150° and 180°, and the θ2 angles also obeys their theoretic values by averaging around 120°.

Note that the θ1 angle in C-X*⋯*Π halogen bonds is on average 127°, which is substantially smaller than the expected 150° till 180° range (Figure2B). This trend is independent of the type of halogen atom.

As no restraints were applied on the θ1 and θ2 angles in the PDB-REDO model generation, these angles are unbiased. The Van der Waals overlap is weakly restrained compared to the distances, so the bias for the Van der Waals overlap is considered to be minimal. Furthermore, our data quality is sufficient for analysis of distributions (Supplemental Figures S2 and S3). Therefore, we proceed with defining target values for all halogen bond categories, except C-I*⋯*Π halogen bonds involving tryptophan, as there are fewer than five observations for these cases. We define a way to judge the quality of halogen-acceptor pairs, while considering that the distributions of the metrics regarding the geometry are mono-disperse, but not normally distributed (Supplemental Figures S4 and S5). Hence, it is not prudent to determine targets as means with standard deviations for a halogen bond.

We evaluate the geometric quality of a potential halogen bond by calculating a score for each relevant geometric parameter, referred to as *Score*_*geom*_, using the following equation:

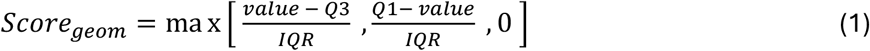

Here, *value* is the measured value of a geometric parameter; *Q1* and *Q3* are the first and third quartiles, respectively; and *IQR* is the interquartile range (Q3 − Q1), derived from the boxplot statistics of the applicable reference dataset. The score expresses the distance of a value from the boundaries of the ‘box’ of the reference dataset in terms of the number of IQRs. Taken that a value within Q1–Q3 (i.e., the interquartile range) is considered preferred, a value within 1.5 × IQR of the quartiles (i.e., within the whiskers) is allowed, and a value outside the whiskers is treated as an outlier, this means that any outlier will have a *Score*_*geom*_ > 1.5

We then apply Equation (1) independently to each geometric parameter relevant to the halogen bond geometry. For C–X··Y interactions, *Score*_*geom*_ is computed for distance, θ1 and θ2; and for C– X⋯Π interactions, it is computed for distance and θ1 only, as θ2 is not defined in this context. We then define an overall Halogen Bond Score (HalBS) as the maximum of the individual geometric scores:

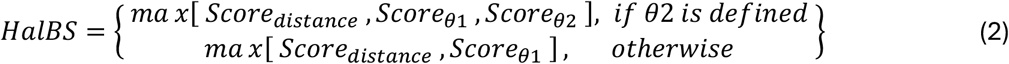

This score reflects the least favourable geometric feature of the halogen bond and is used as a proxy for assessing bond quality. A HalBS > 1.5 indicates that at least one geometric parameter is an outlier, suggesting that the halogen bond geometry may be abnormal and should be examined more closely.

The robustness and accuracy of HalBS improve with the quantity and diversity of data used to compute the reference boxplot statistics. For example, in C–X⋯Π interactions involving amino acids like tyrosine, histidine, or tryptophan, limited available data may affect the reliability of the scoring.

### HalBS implementation in PDB-REDO

A reference implementation of HalBS reports scores for C-X··O, C-X··S, C-X*⋯*His, C-X*⋯*Phe, C-X*⋯*Tyr and C-X*⋯*Trp halogen bonds where X=F, Cl, Br, I, with the exception of C-I*⋯*Trp, for which boxplot statistics could not be calculated due to insufficient data (Supplemental Table S4). Halogen atoms in alternate conformations are excluded, as well as water molecules functioning as potential acceptor. Furthermore, the θ1 angle between the halogen atom and its potential acceptor must be larger than 90°. When multiple halogen bonds are possible for a single halogen atom (C-X··Y and C-X*⋯*Π combined), the bond with the lowest HalBS score is selected and reported. Note that while the boxplots were generated for halogen containing ligands, the implementation of HalBS is calculated for any type of halogen containing compound, including non-canonical amino acids and nucleotides. The implementation is added to PDB-REDO (version 8.16) which now reports the number of halogen bonds and the mean HalBS for each structure model in mmCIF (Westbrook et al. 2022) format. Hereby we aim to increase awareness and create more data to improve the halogen bond definitions and validation pipelines in the future. PDB-REDO entries with detected halogen bonds can be retrieved through the PDB-REDO archive manager at https://pdb-redo.eu/archive/.

### Example cases

The binding of the anti-microbial agent triclosan (TCS) to its target protein enoyl-[acyl-carrier-protein] reductase (ENR), one of the proteins involved in fatty acid synthesis in bacteria, causes ENR to change to a closed conformational state, as seen in PDB entry 3GR6 (Priyadarshi et al. 2010). Besides the described electrostatic interactions between the binding pocket and triclosan, the inhibitor forms a halogen bond with its chlorine close to the carbonyl oxygen atom of the Ala97 backbone (Figure 3A). Notably, this alanine is in the loop that is involved in the conformation change suggesting that the halogen bond contributes to the change in conformation of the loop in ENR.

**Figure 3.**
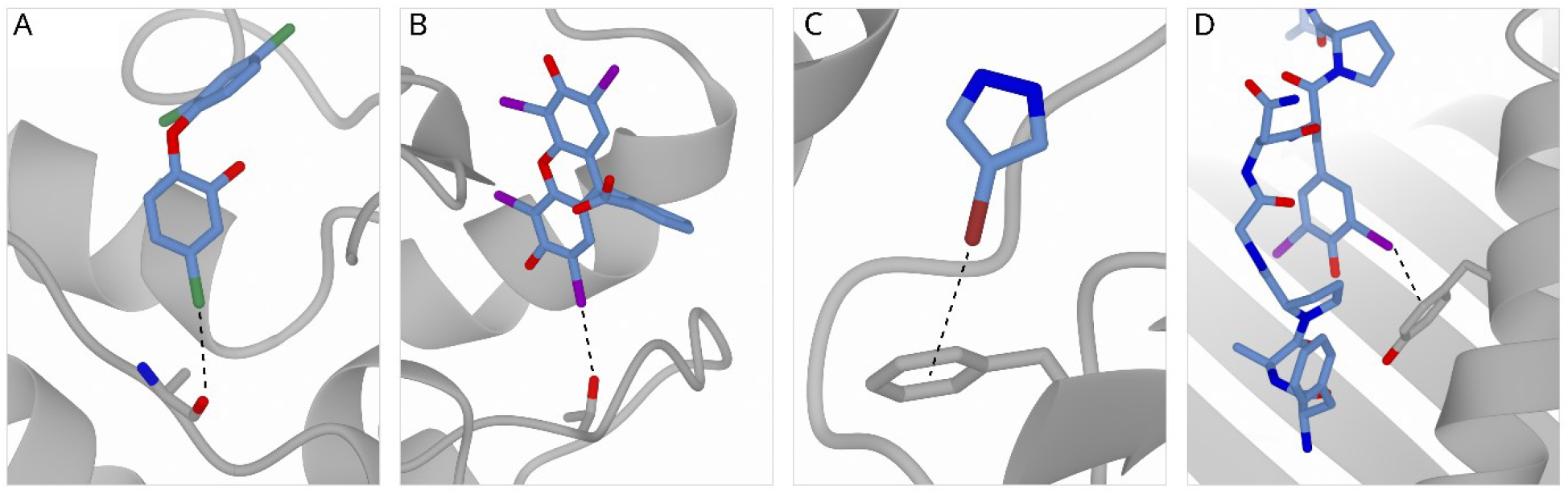
Examples of halogen bonds (marked as a black dotted lines). A) C-X··Y halogen bond between chlorine of triclosan (atom CL14 of TCL A371) and the carbonyl oxygen atom of alanine A97 (PDB-ID: 3GR6, HalBS: 0.36. B) C-X··Y halogen bond between iodine of erythrosine extra bluish (atom I of 9ZZ A303) and hydroxylic oxygen of threonine A77 (PDB-ID 5OOH, HalBS: 1.63. C) C-X⋯Π halogen bond between bromine atom BR4 of BYZ B201 with the π-system of phenylalanine B154 (PDB-ID: 8CZM, HalBS: 0.07) D) C-X⋯Π halogen bond between iodine (atom I1 of TYI C6) with the π-system of tyrosine A159 (PDB-ID 4PGC, HalBS: 0.90).

An example where halogen bonds play a role is in drug development is erythrosin extra bluish, which was found to be a hit compound in a study targeting biliverdin reductases type B (BLVRB) (Nesbitt et al. 2018). In PDB entry 5OOH we observe that the compound has multiple interactions with the protein, among which the iodine atom, that is forming a C-X··Y halogen bond (Figure 3B).

Halogen atoms often appear in crystallographic fragment screening experiments, where multiple small organic compounds are screened towards a target protein. Bacterial oxidoreductase in *E*.

*coli* DsbA (EcDsbA) is such a target. It is responsible for the formation of disulfide bonds into many bacterial factors causing disease, and hence is considered a target in the treatment of drug resistant bacterial infections (Whitehouse et al. 2023). In PDB entry 8CZM, one of the bromine-containing fragments is BYZ (4-bromo-1H-pyrazole), which binds to EcDsbA and forms a C-X*⋯*Π halogen bond with the π-system of phenylalanine in the protein (Figure 3C).

An example of a modified amino acid containing halogens is 3,5-diiodotyrosine (TYI), which was used to investigate peptide selectivity in the major histocompatibility complex (MHC) class I (PDB-ID 4PGC) (Garstka et al. 2015). In this study, increasingly larger residues were used to probe the peptide binding site of MHC class I. Although the iodine atoms were used for their size, one iodine makes a clear C-X*⋯*Π halogen bond with a tyrosine in the binding site (Figure 3D).

The examples described above demonstrate perfectly fine halogen bonds according to theory. The halogen bonds involving triclosan, the BYZ fragment and the modified amino acid TYI (Figures 3A, C and D, respectively) all have a HalBS <1.5. indicating a preferred halogen bond geometry. However, the halogen bond involving erythrosine extra bluish (Figure 3B) adopts a score >1.5, suggesting it should be considered with caution. We suggest that the current HalBS cannot be used as a direct validation metric but can provide indication of genuine halogen bonds and “not so proper” halogen bonds.

### Future considerations

To further optimize validation of C-X··Y halogen bonds it could be investigated whether different types of halogen bond geometries are possible (Ibrahim et al. 2022). Separating C-X··O halogen bonds into carbonyl oxygen, carboxyl and hydroxyl oxygen atoms might improve the accuracy of halogen bond descriptions (Supplemental Figure S6) (Costa 2017; Zhang et al. 2017). Another challenge is to identify C-X··N halogen bonds with histidine as acceptor. The histidine protonation state defines whether such interaction is possible, but depending of the local molecular context reliably establishing this can range from straightforward to computationally expensive (Hooft et al. 1996; Evans and Murshudov 2013). It should also be considered to relate our findings to halogen bonds identified by Arpeggio (Jubb et al. 2017), which uses slightly different definitions for halogen bond distances and angles.

To further investigate C-X*⋯*Π halogen bonds, the distance from the halogen perpendicular to the π-system of the acceptor is worth to calculate and include as geometric parameter (Riley and Tran 2017). Another option could be to use a θ2-equivalent angle between the halogen atom: the centroid of the π-system and one carbon atom in the π-system, which has an ideal value around 90°. This should ease the filtering of halogen-acceptor pairs into actual halogen bonds wherein the σ-hole of the halogen is positioned perpendicular to the π-system.

More critical filtering or more accurate data can distinguish between a genuine halogen bond and “a halogen in proximity of a possible acceptor atom”, which will be particularly effective for fluorine and chlorine. To add more observations and hereby increase the dataset size, halogen bonds between other types of macromolecules can be considered, like modified amino acids or nucleotides. Also, halogen bonds can be formed with the π-system of peptide bonds (Auffinger et al. 2004). Identifying such interactions in PDB-REDO structure models would add an extra halogen bond type to the dataset that we have not examined here.

Additionally, interactions between halogen atoms and water molecules could be investigated further. While visually inspecting hundreds of example cases, we observed false positives where the halogen bond with a water molecule would be more likely than the acceptor we found while datamining. These should be considered in future pipelines. Note that with water molecules there comes an extra complication, as the location of the free lone pairs depends on the hydrogen bonding network and is not always clear.

Besides these optimizations with respect to defining halogen-acceptor pairs, more (high quality) data could contribute to the differentiation between a genuine halogen bond and “a halogen in proximity of a possible acceptor”. A valuable source of potential ligand-protein halogen bonds can be crystallographic fragment screening data. The fragment libraries used in such screens typically hold many halogen containing compounds, albeit more fluorine and chlorine than bromine and iodine atoms. Overcoming data archiving issues (Jaskolski et al. 2022; Weiss et al. 2022; Erlanson et al. 2025; Perrakis 2025) could allow valuable data from hundreds or even thousands of ligand-protein complexes obtained from fragment screening to enrich halogen bond observations and hereby ease the definition of validation targets.

## Conclusion

We identified halogen bonds between ligands and proteins in PDB-REDO structure models. With HalBS, we introduce a metric that annotates halogen bonds adopting preferred or allowed geometry, or are an outlier and should be examined critically. This approach is integrated into the PDB-REDO pipeline and can be used in other ligand validation pipelines. By increasing awareness and accumulating (high quality) data of halogen bonds in ligand-protein complexes, it could become possible to determine validation targets with ideal values and corresponding standard deviations for each halogen-acceptor pair.

## Methods

### Data mining

Compound IDs of halogen containing compounds were mined from the Chemical Component Dictionary (CCD, (Westbrook et al. 2015)) of the Protein Data Bank (Burley et al. 2019) and stored with their type and the name(s) of the halogen atom(s) in the molecule. Compounds in which the halogen atom is not connected to a carbon atom, e.g. in halogen-metal complexes and halogen acids, and the single ion halogens (F-, Cl-, Br-, I-) were removed from this list to focus on organic halogen bonds. The identifiers of excluded halogen containing compounds are: F, CL, BR, IOD, 08T, 0JC, 0OD, 0TE, 202, 2T8, 4IR, 4KV, 5LN, 61C, 61D, 6BP, 6O0, 73M, 7GE, 8TH, 8TR, 8WV, 9QB, 9RU, 9TH, A9J, AF3, ALF, BE7, BEF, BF2, BF4, BFD, BPT, C2C, C7P, CFO, CPT, CUL, D0X, D7Z, DAA, DAE, DAQ, E3D, E5O, ELJ, F6Q, F7T, FPO, HG2, HGI, I2I, I3M, I83, J0K, J0N, JR3, KQB, KYS, KYT, LCO, LCP, LN8, MF4, MGF, MNQ, N2N, N2R, N2S, N2W, NG8, NMQ, NXC, O1N, ONP, ORS, OS1, OT1, P3C, PC4, PCL, PEJ, PNQ, PT7, QLT, R1N, RAX, RBN, RHE, RSW, RU0, RU7, RUD, RUH, SFL, SRX, SVP, SXC, TBR, TPT, U0J, VKZ, VL2, YPT, YXX, YXZ, ZN0, ZN5, ZN6, ZN7, ZN8, ZN9, ZPT.

### Defining halogen bonds in PDB-REDO structure models

For each of the PDB-REDO (Joosten et al. 2009) structure models with at least one halogen containing compound, the halogen atoms were mined and stored with their possible acceptor atoms. An acceptor atom is an oxygen (O), nitrogen (N), sulfur (S), phosphor (P) or selenium (Se) atom in an organic compound within 4Å of the halogen atom itself. For each halogen-acceptor pair, the distance between the two atoms and their Van der Waals overlap, as well as the θ1 and θ2 angles were calculated. The Van der Waals radii of the atoms were taken from https://en.m.wikipedia.org/wiki/Atomic_radii_of_the_elements_(data_page) (Bondi 1964; Slater 1964; Clementi et al. 1967; Pyykkö et al. 2005; Mantina et al. 2009). Possible halogen bonds to water were excluded as θ2 angles could not be calculated.

Similarly, as for C-X··Y halogen bonds, we mined halogen bonds between a halogen atom and a π-system found in either phenylalanine (PHE), tyrosine (TYR), histidine (HIS) or tryptophan (TRP) to find halogen-π bonds within protein-ligand complexes. Such an interaction was assigned when the distance between the halogen atom and the centroid of the π-system was within 6Å. The centroid of the π-system was also used to calculate the θ1 angle for the interaction. For the halogen-π interactions, θ2 and the Van der Waals overlap cannot be calculated.

When a halogen atom had multiple possible acceptors, the acceptor with the θ1 angle closest to 180° was selected for further analyses to get a single acceptor for each halogen atom. Halogen-acceptor pairs with a θ1 angle <90° were removed from the data, as such angles would render the σ-hole unreachable (Figure 1).

### Mining metadata

For each structure model in which we found a halogen bond, the resolution of the X-ray data, the R-factor, the R-free, average B-factor and PDB-REDO version were stored. Finally, the real space correlation coefficient (RSCC, (Jones et al. 1991)), as calculated by density-fitness (van Beusekom et al. 2019), was mined for both the halogen containing compound as well as the compound containing the acceptor.

### Software

Mining of the halogen bonds was done using libcif++ (Hekkelman) and libpdb-redo (Hekkelman). The obtained datasets were loaded using pandas2.1.4 (The pandas development team 2020), plots were made using Seaborn0.13.2 (Waskom 2021), all in Python3.12.1.

## Supplemental material

Supplemental material figures and tables referenced in this manuscript are available in PDF format.

## Data availability statement

Reference code for the implementation of HalBS score calculations in PDB-REDO is available at https://github.com/PDB-REDO/HalBS.

## Acknowledgements and funding

We thank the Research High Performance Computing facility of the Netherlands Cancer Institute for providing and maintaining computation resources. This project was funded by the Horizon2020 EC projects iNEXT-Discovery (871037) and FragmentScreen (101094131), and by an institutional grant of the Dutch Cancer Society and of the Dutch Ministry of Health, Welfare and Sport. We also thank Oncode Institute and CCP4 for their financial support and Pantelis G. Bagos for fruitful discussions on the data analysis.

