## Supplemental information for "Halogen bonds between ligands and proteins: can we use them in validation?"

**Supplemental Figure S1** Overall distributions of the halogen bonds in PDB-REDO, from left to right: Resolution of the structures wherein halogen bonds were found, B-factors of halogen atoms and their acceptors, RSCC (real space correlation coefficient) of donor and acceptors residues. Bin-widths from left to right: 0.1,10,10,0.01,0.01. A) quality indicators for C-X· ·Y bonds; B) quality indicators for C-X· · ·Π bonds.

**A**

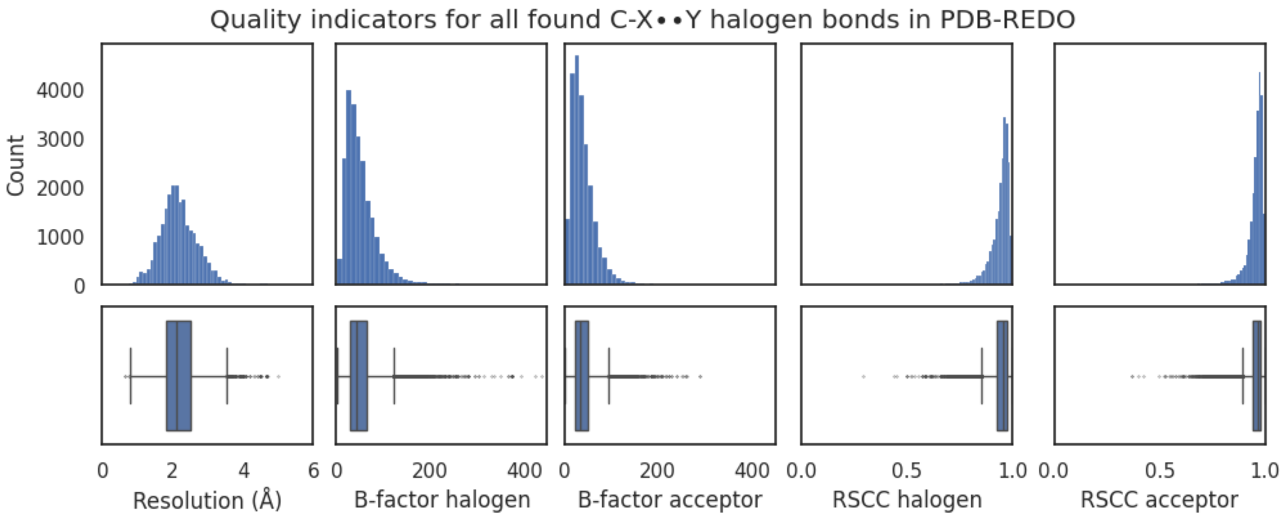


**B**

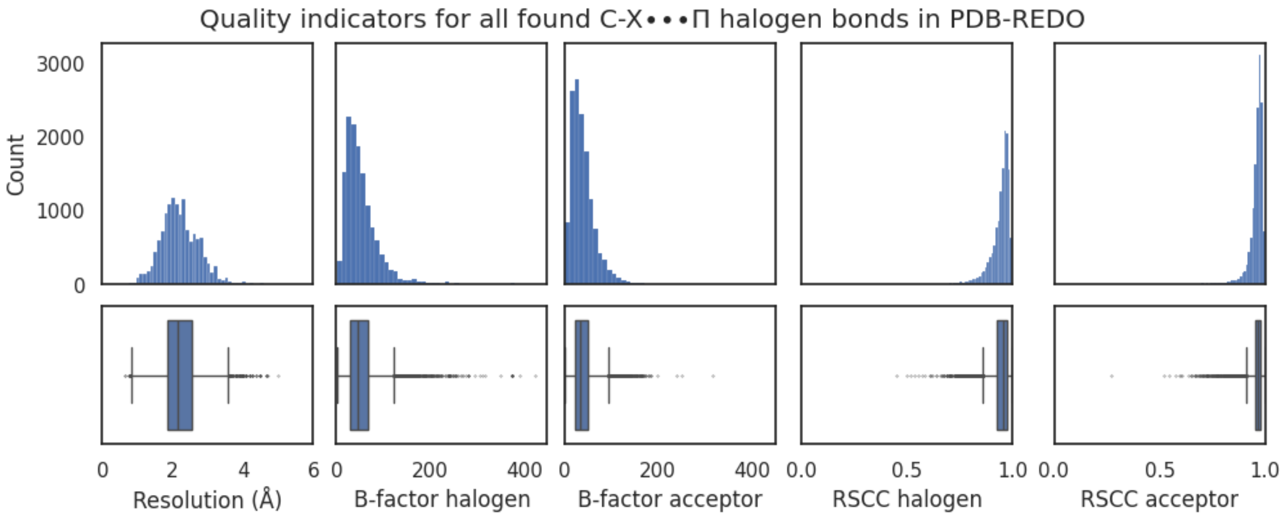

**Supplemental Figure S2** Distributions of the quality indicators for the ligand-protein C-X· ·Y halogen bonds found in PDB-REDO after initial filtering. From left to right: Resolution of the structures wherein halogen bonds were found, B-factors of halogen atoms and their acceptors, RSCC (real space correlation coefficient) of the compounds containing the halogen or the acceptor atom. Bin-widths from left to right: 0.1,10,10,0.01,0.01. A) quality indicators for C-X· ·Y bonds with fluorine; B) quality indicators for C-X· ·Y bonds with chlorine; C) quality indicators for C-X· ·Y bonds with bromine; D) quality indicators for C-X· ·Y bonds with iodine.

**A**

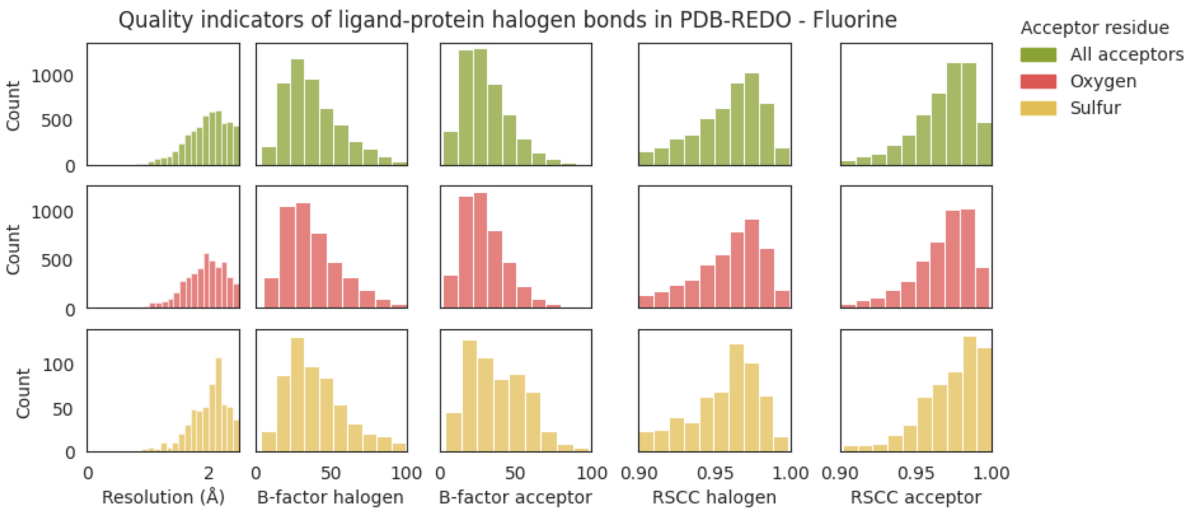


**B**

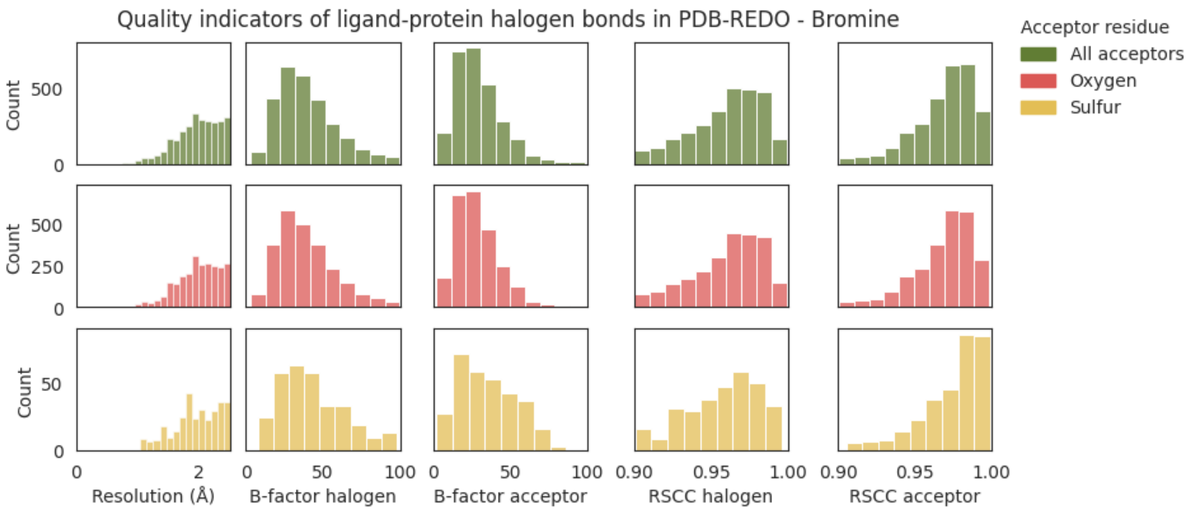

**C**

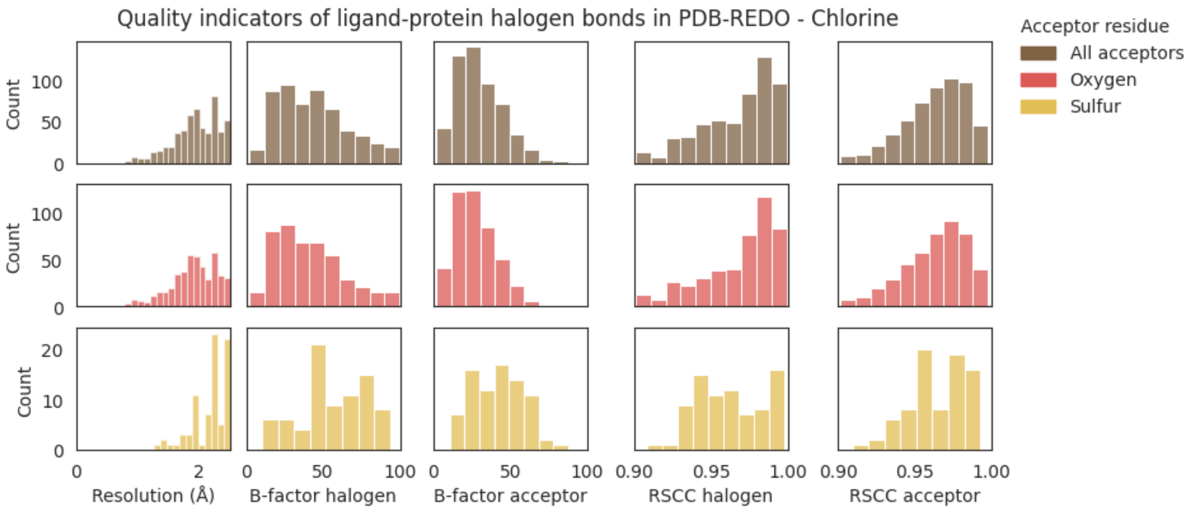

Quality indicators of ligand-protein halogen bonds in PDB-REDO - Chlorine
Acceptor residue
All acceptors
Oxygen
Sulfur
Count
Resolution (Å)
B-factor halogen
B-factor acceptor
RSCC halogen
RSCC acceptor


**D**

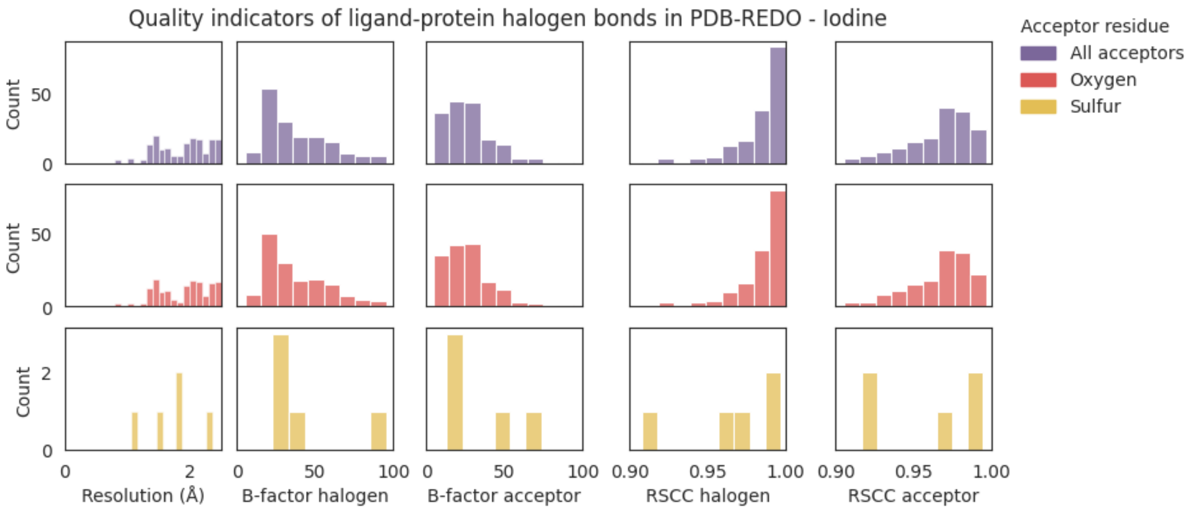

Quality indicators of ligand-protein halogen bonds in PDB-REDO - Iodine
Acceptor residue
All acceptors
Oxygen
Sulfur
Count
Resolution (Å)
B-factor halogen
B-factor acceptor
RSCC halogen
RSCC acceptor

**Supplemental Figure S3** Distributions of the quality indicators for the ligand-protein C-X···Π halogen bonds found in PDB-REDO after initial filtering. From left to right: Resolution of the structures wherein halogen bonds were found, B-factors of halogen atoms and their acceptors, RSCC (real space correlation coefficient) of the donor and acceptor residues. Bin-widths from left to right: 0.1,10,10,0.01,0.01. A) quality indicators for C-X···Π bonds with fluorine; B) quality indicators for C-X···Π bonds with chlorine; C) quality indicators for C-X···Π bonds with bromine; D) quality indicators for C-X···Π bonds with iodine.

**A**

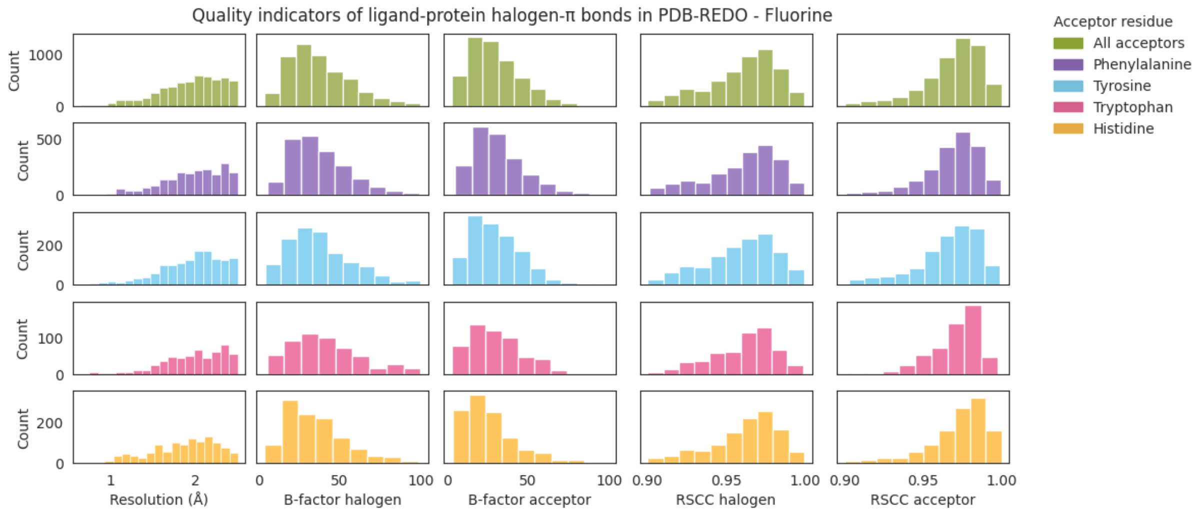


**B**

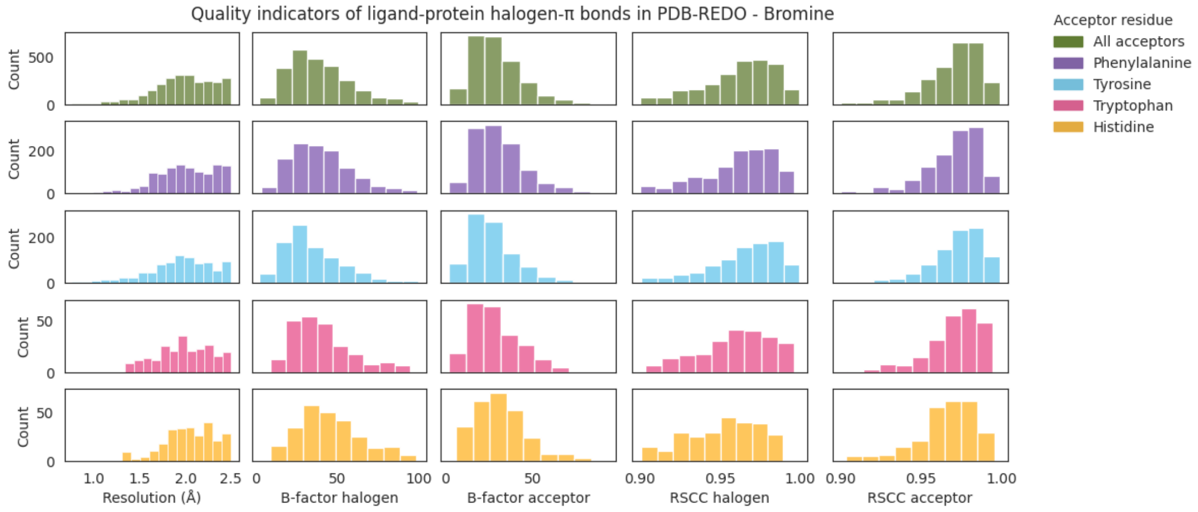

**C**

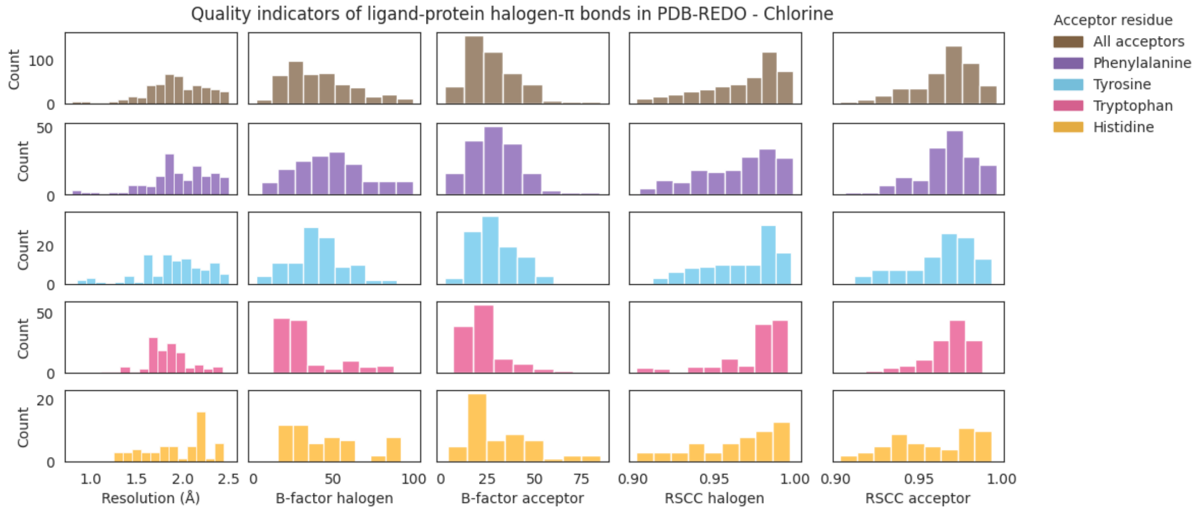

Quality indicators of ligand-protein halogen-π bonds in PDB-REDO - Chlorine
Count
Resolution (Å)
B-factor halogen
B-factor acceptor
RSCC halogen
RSCC acceptor
Acceptor residue
All acceptors
Phenylalanine
Tyrosine
Tryptophan
Histidine


**D**

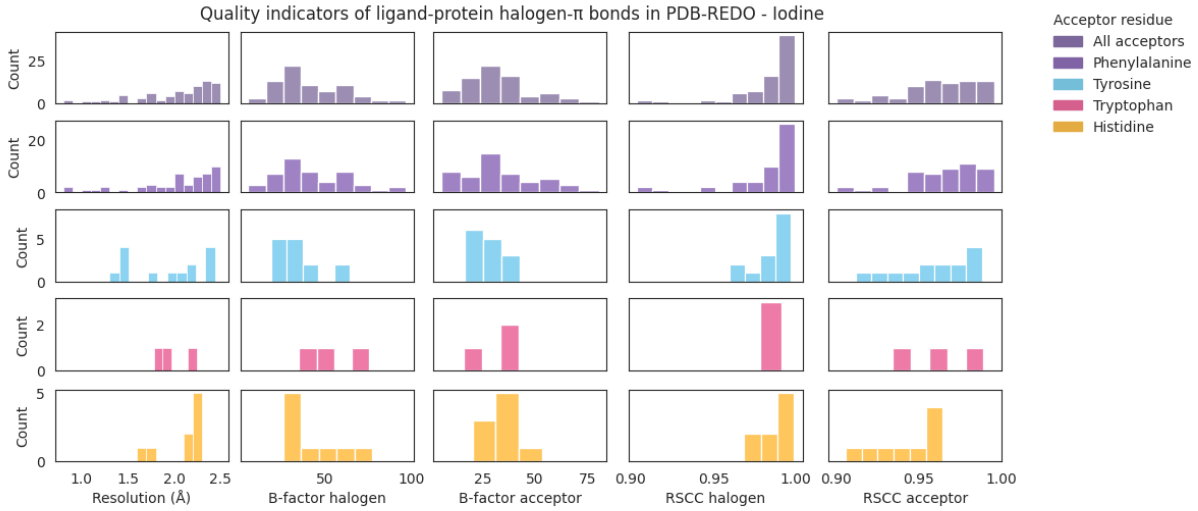

Quality indicators of ligand-protein halogen-π bonds in PDB-REDO - Iodine
Count
Resolution (Å)
B-factor halogen
B-factor acceptor
RSCC halogen
RSCC acceptor
Acceptor residue
All acceptors
Phenylalanine
Tyrosine
Tryptophan
Histidine

**Supplemental Figure S4** Distributions of the geometric parameters for the ligand-protein C-X··Y halogen bonds found in PDB-REDO. From top to bottom: halogen bonds containing fluorine, chlorine, bromine and iodine, respectively. From left to right: All halogen bonds, halogen bonds with acceptor O-C, N-C, S-C, respectively. Y-axis show counts. A) Distribution of distances, bin-width 0.1A. B) Distribution of van-der-Waals overlap, bin-width 0.1Å. C) Distribution of θ1 angles, bin-width 5˚. D) Distribution of θ1 angles, bin-width 5˚.

**A**

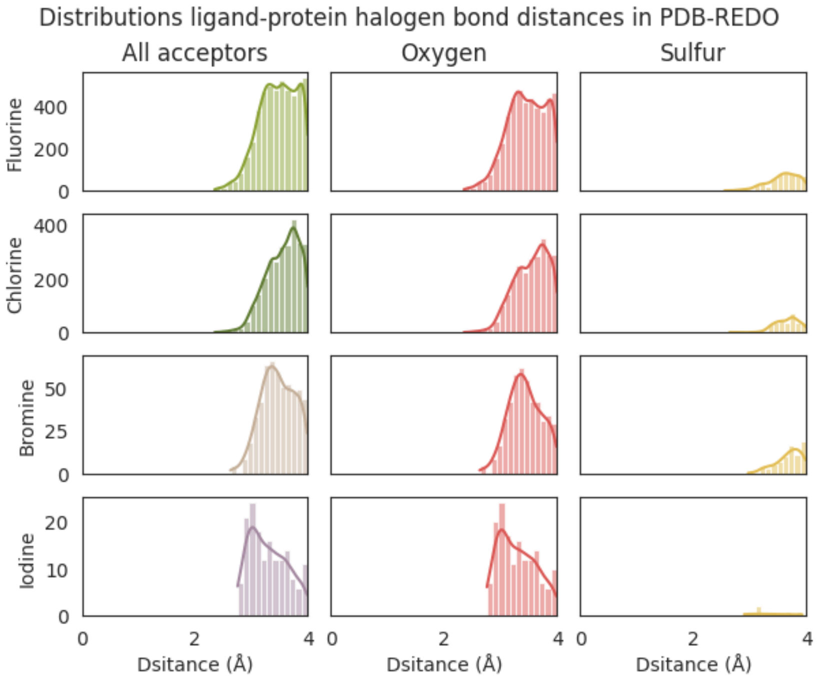


**B**

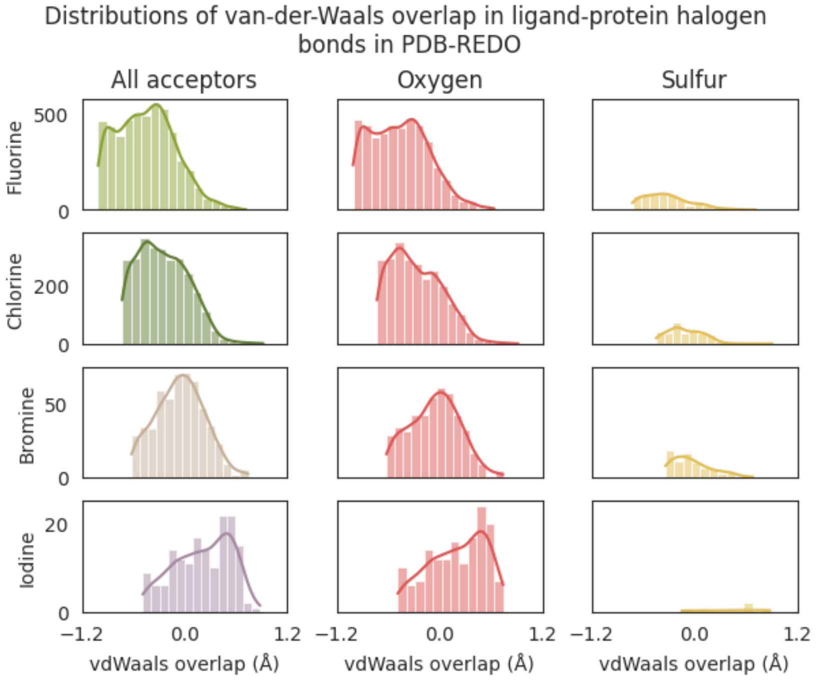

C

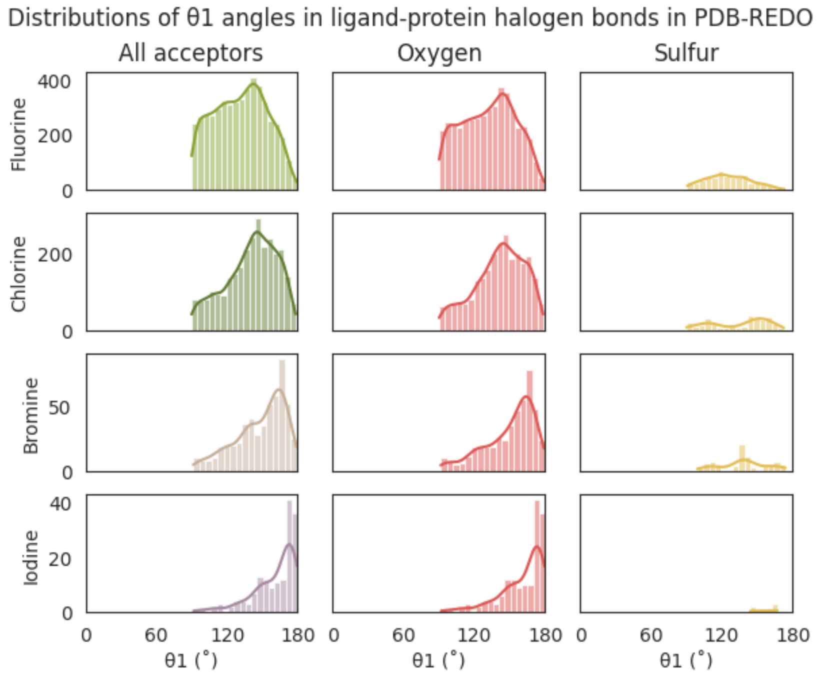
Distributions of θ1 angles in ligand-protein halogen bonds in PDB-REDO
All acceptors
Oxygen
Sulfur
Fluorine
Chlorine
Bromine
Iodine
400
200
0
50
40
20
0 60 120 180
θ1 (°)

D

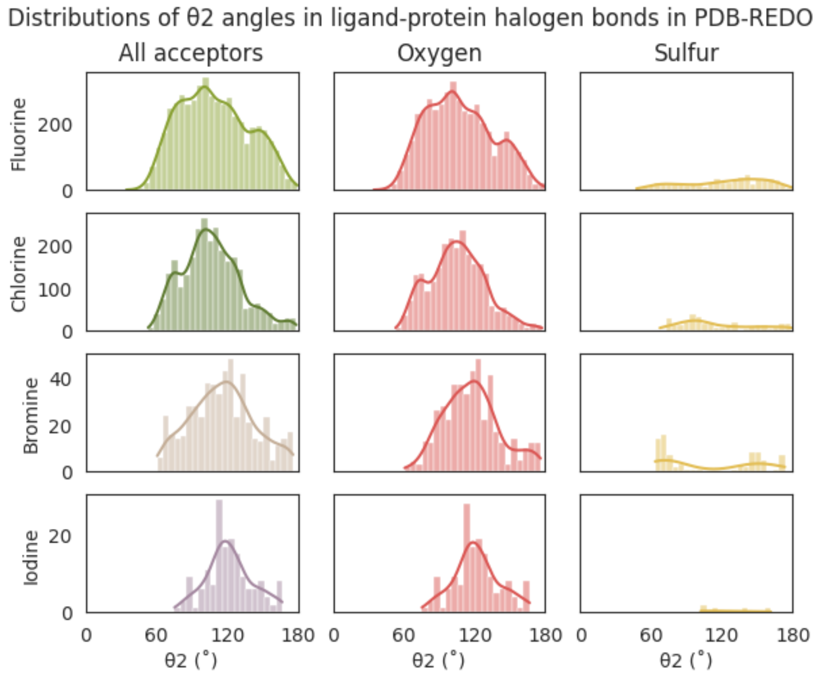
Distributions of θ2 angles in ligand-protein halogen bonds in PDB-REDO
All acceptors
Oxygen
Sulfur
Fluorine
Chlorine
Bromine
Iodine
200
100
0
40
20
0 60 120 180
θ2 (°)

**Supplemental Figure S5** Distributions of the geometric parameters for the ligand-protein C-X···Π halogen bonds found in PDB-REDO. From top to bottom: halogen bonds containing fluorine, chlorine, bromine and iodine, respectively. From left to right: All halogen bonds, halogen bonds with acceptor phenylalanine, tyrosine, tryptophan and histidine, respectively. Y-axis show counts. A) Distribution of distances, bin-width 0.1Å. B) Distribution of θ1 angles, bin-width 5˚.

**A**

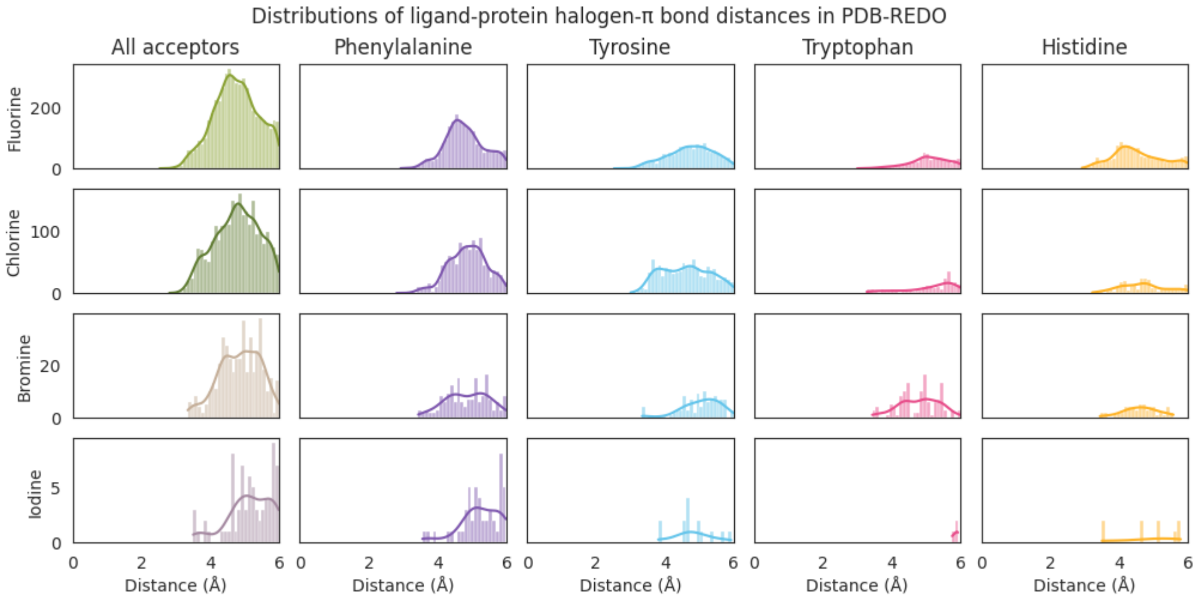


**B**

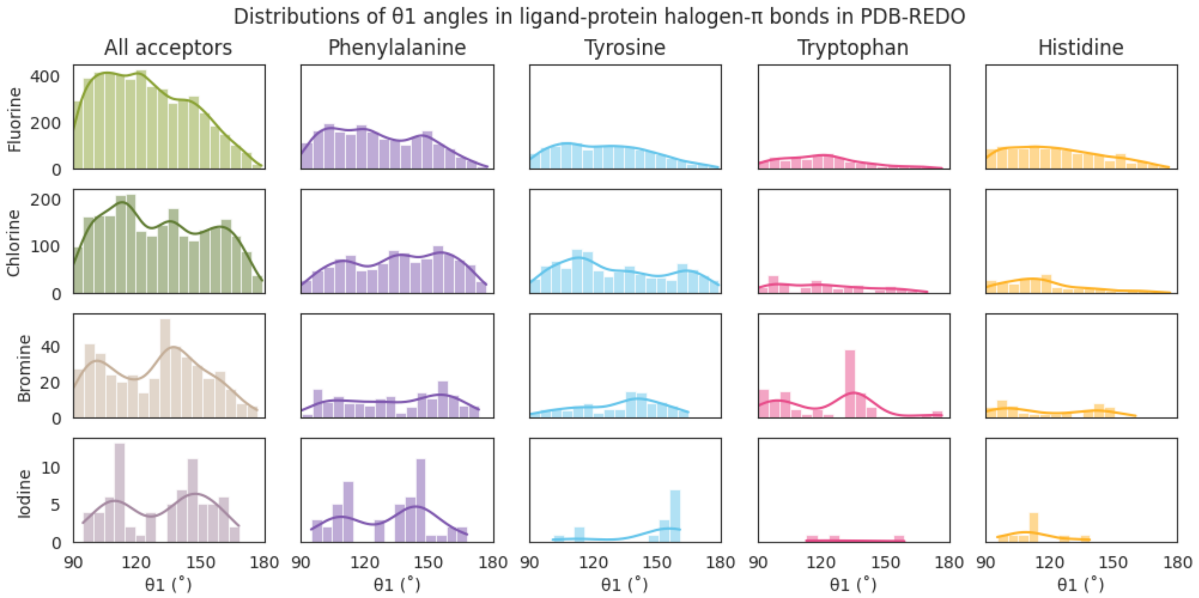

**Supplemental Figure S6** Geometry of ligand-protein C-X· ·O halogen bonds found in PDB-REDO. For each halogen (from left to right: fluorine (light green), chlorine (dark green), bromine (brown), iodine (purple)) the overall distance, van-der-Waals overlap, θ1 and θ2 angles are shown on the left panel. In the panels on the right-hand side these values are split by acceptor oxygen type: Hydroxyl, Carboxyl and Carbonyl. Note that a positive value for the sum of the van-der-Waals radii means that the distance between halogen and acceptor atom is shorter than the van-der-Waals overlap. For clarity a strip plot is shown when there are 25 or fewer observations.

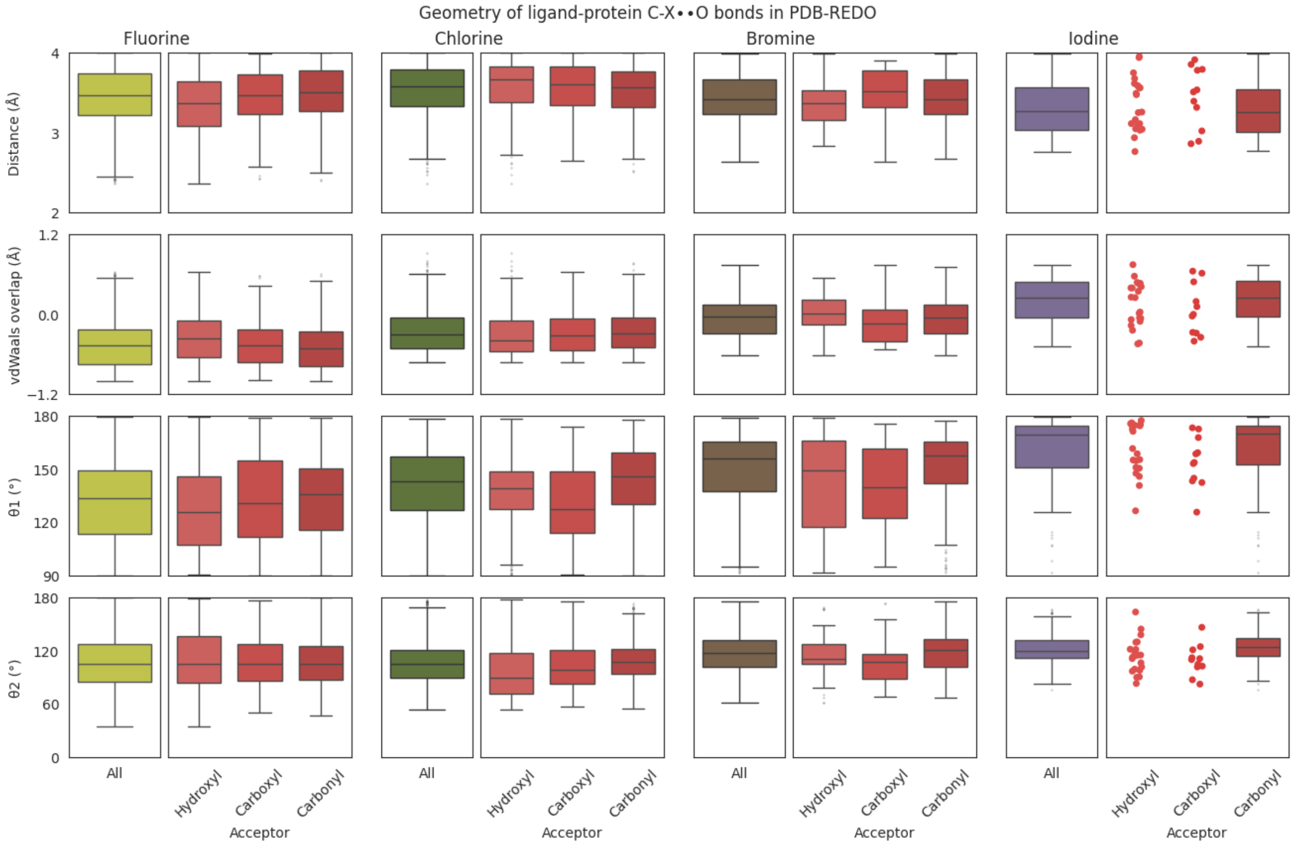

**Supplemental Table S1** Number of observations of halogen-acceptor pairs for C-X··Y and C-X···Π bonds found in PDB-REDO structure models, with a crystallographic resolution of 2.5Å or better, a B-factor of 100 $Å^2$ or lower, and an RSCC equal to or greater than 0.9 for the donating and accepting residues. For C-X··Y bonds the acceptor is listed with its neighbouring atom, e.g. O-C is an oxygen atom as acceptor, which is connected to a carbon atom. For the accepting π-system is noted as the amino acid wherein it was found, e.g. Phe indicates that there is a halogen bond between the halogen and the π-system of the phenyl ring in the phenylalanine side chain.

| | Number of halogen-acceptor pairs | | | | |
|---|---|---|---|---|---|
| Halogen / Acceptor | F | Cl | Br | I | Total |
| | **C-X··Y halogen bonds** | | | | |
| **O-C** | 4640 | 2548 | 515 | 172 | **7875** |
| **S-C** | 571 | 314 | 80 | 11 | **976** |
| **S-S** | 3 | 4 | 0 | 1 | **8** |
| **O-P** | 3 | 1 | 1 | 0 | **5** |
| **O-O** | 1 | 0 | 1 | 0 | **2** |
| **O-S** | 1 | 0 | 0 | 0 | **1** |
| **P-O** | 2 | 0 | 0 | 0 | **2** |
| **Se-C** | 0 | 0 | 0 | 1 | **1** |
| **Total** | **5221** | **2867** | **597** | **185** | **8870** |
| | **C-X···Π bonds** | | | | |
| **Phe** | 2146 | 170 | 1114 | 51 | **3481** |
| **Tyr** | 1389 | 125 | 908 | 20 | **2442** |
| **His** | 1216 | 63 | 258 | 12 | **1549** |
| **Trp** | 584 | 122 | 226 | 3 | **935** |
| **Total** | **5335** | **2506** | **480** | **86** | **8407** |

**Supplemental Table S2** The compound types found in the CCD (Chemical Component Dictionary) of the halogen atom containing compounds for the observed halogen bonds.

| | **Number of compounds** | |
|---|---|---|
| **Compound type** | **C-X· · ·Y** (Y = O-C or S-C) | **C-X· · ·Π** |
| Non-polymer | 8442 | 8096 |
| l-peptide linking | 153 | 64 |
| d-peptidelinking | 17 | 8 |
| Peptide-like | 13 | 18 |
| DNA-linking | 79 | 51 |
| RNA-linking | 25 | 21 |
| l-saccharide | 1 | 3 |
| l-saccharide-alphalinking | 8 | 5 |
| l-saccharide-betalinking | 0 | 3 |
| d-saccharide | 30 | 40 |
| d-saccharide-alphalinking | 46 | 64 |
| d-saccharide-betalinking | 37 | 34 |

**Supplemental Table S3** Statistical values (number of observations, mean, median, Q1, Q3 and Median absolute deviation (MAD)) for the distance, Van-der-Waals overlap, θ1 and θ2 angles in C-X· ·Y halogen bonds corresponding to the boxplots shown in Figure 2A.

| | **F all** | **F· ·O** | **F· ·S** | **Cl all** | **Cl· ·O** | **C· ·S** | **Br all** | **Br· ·O** | **Br· ·S** | **I all** | **I· ·O** | **I· ·S** |
|---|---|---|---|---|---|---|---|---|---|---|---|---|
| **Count** | 4935 | 4381 | 554 | 2791 | 2479 | 312 | 536 | 456 | 80 | 161 | 156 | 5 |
| **Distance (Å)** | | | | | | | | | | | | |
| **Mean** | 3.5 | 3.5 | 3.6 | 3.6 | 3.5 | 3.7 | 3.5 | 3.4 | 3.7 | 3.3 | 3.3 | 3.4 |
| **Median** | 3.5 | 3.5 | 3.6 | 3.6 | 3.6 | 3.7 | 3.5 | 3.4 | 3.7 | 3.3 | 3.3 | 3.2 |
| **Q1** | 3.2 | 3.2 | 3.4 | 3.4 | 3.3 | 3.5 | 3.3 | 3.2 | 3.6 | 3.0 | 3.0 | 3.1 |
| **Q3** | 3.8 | 3.7 | 3.8 | 3.8 | 3.8 | 3.8 | 3.7 | 3.7 | 3.9 | 3.6 | 3.6 | 3.7 |
| **MAD** | 0.3 | 0.3 | 0.2 | 0.2 | 0.2 | 0.1 | 0.2 | 0.2 | 0.2 | 0.3 | 0.3 | 0.3 |
| **Van-der-Waals overlap (Å)** | | | | | | | | | | | | |
| **Mean** | -0.5 | -0.5 | -0.3 | -0.3 | -0.3 | -0.1 | -0.1 | -0.1 | 0.0 | 0.2 | 0.2 | 0.4 |
| **Median** | -0.5 | -0.5 | -0.4 | -0.3 | -0.3 | -0.1 | -0.1 | 0 | -0.1 | 0.2 | 0.2 | 0.6 |
| **Q1** | -0.7 | -0.8 | -0.5 | -0.5 | -0.5 | -0.3 | -0.3 | -0.3 | -0.2 | -0.1 | -0.1 | 0.1 |
| **Q3** | -0.2 | -0.2 | -0.2 | 0.0 | -0.1 | 0.0 | 0.1 | 0.1 | 0.1 | 0.5 | 0.5 | 0.6 |
| **MAD** | 0.2 | 0.3 | 0.2 | 0.2 | 0.2 | 0.1 | 0.2 | 0.2 | 0.2 | 0.3 | 0.3 | 0.3 |
| **θ1 (˚)** | | | | | | | | | | | | |
| **Mean** | 131.2 | 132.0 | 125.2 | 140.0 | 140.8 | 134.3 | 147.8 | 149.5 | 138.6 | 160.5 | 160.6 | 157.9 |
| **Median** | 132.1 | 133.4 | 123.5 | 143.2 | 143.2 | 143.2 | 154.0 | 155.7 | 137.7 | 169.2 | 169.6 | 164.4 |
| **Q1** | 112.9 | 113.1 | 111.1 | 125.4 | 126.8 | 110.4 | 135.9 | 137.2 | 131.2 | 150.4 | 150.8 | 148.4 |
| **Q3** | 148.5 | 149.5 | 138.5 | 156.9 | 157.3 | 155.4 | 165.0 | 165.4 | 154.3 | 174.0 | 174.4 | 165.6 |
| **MAD** | 17.6 | 17.9 | 14 | 15.2 | 15 | 16.6 | 13.6 | 12.2 | 13.2 | 6.9 | 7 | 2.5 |
| **θ2 (˚)** | | | | | | | | | | | | |
| **Mean** | 109.4 | 107.8 | 122 | 106.7 | 105.5 | 116.3 | 116.4 | 117.9 | 107.9 | 122.0 | 122 | 123.9 |
| **Median** | 106.5 | 104.8 | 129.4 | 104.7 | 104.7 | 105.4 | 117.4 | 117.8 | 84.7 | 120.0 | 120.1 | 114.2 |
| **Q1** | 85.7 | 85.3 | 95.1 | 89.8 | 89.2 | 93.2 | 97.2 | 101.3 | 69.9 | 111.7 | 111.9 | 105.1 |
| **Q3** | 130.7 | 127.3 | 151.5 | 122.4 | 121.1 | 145 | 133.5 | 132.2 | 150.2 | 131.6 | 131.6 | 136.9 |
| **MAD** | 22.4 | 20.8 | 25.4 | 16.3 | 16.1 | 21.7 | 17.9 | 15.2 | 18.6 | 10.1 | 9.7 | 12.2 |

**Supplemental Table S4** Statistical values (number of observations, mean, median, Q1, Q3 and Median absolute deviation (MAD)) for the distance, and θ1 angles in C-X···Π halogen bonds corresponding to the boxplots shown in Figure 2B. Cases with fewer than 5 observations are marked as 'NA' as no boxplot statistics can be calculated.

| | **F all** | **F···Phe** | **F···Tyr** | **F···Trp** | **F···His** | **Cl all** | **Cl···Phe** | **Cl···Tyr** | **Cl···Trp** | **Cl···His** |
|---|---|---|---|---|---|---|---|---|---|---|
| **Count** | 5099 | 2097 | 1328 | 539 | 1135 | 2477 | 1100 | 895 | 225 | 257 |
| **Distance (Å)** | | | | | | | | | | |
| **mean** | 4.7 | 4.8 | 4.8 | 5.1 | 4.5 | 4.8 | 4.9 | 4.6 | 5.1 | 4.7 |
| **median** | 4.7 | 4.7 | 4.8 | 5.0 | 4.4 | 4.8 | 4.9 | 4.6 | 5.4 | 4.7 |
| **Q1** | 4.3 | 4.4 | 4.3 | 4.7 | 4.1 | 4.3 | 4.5 | 4.0 | 4.6 | 4.2 |
| **Q3** | 5.2 | 5.1 | 5.2 | 5.5 | 5.0 | 5.3 | 5.3 | 5.2 | 5.7 | 5.1 |
| **MAD** | 0.4 | 0.3 | 0.5 | 0.4 | 0.4 | 0.5 | 0.4 | 0.6 | 0.4 | 0.5 |
| **θ1 (˚)** | | | | | | | | | | |
| **mean** | 124.6 | 125.7 | 125.3 | 120.7 | 123.4 | 130.6 | 135.4 | 130.6 | 121.6 | 117.6 |
| **median** | 122.2 | 123.1 | 123.8 | 119.2 | 121.1 | 129.3 | 137.0 | 126.2 | 119.0 | 114.8 |
| **Q1** | 106.8 | 107.5 | 107.4 | 105.6 | 105.9 | 110.7 | 115.0 | 110.7 | 100.6 | 103.3 |
| **Q3** | 141.0 | 144.4 | 141.1 | 131.6 | 139.0 | 151.2 | 155.0 | 151.8 | 137.1 | 127.4 |
| **MAD** | 16.8 | 18.1 | 16.7 | 13.1 | 16.5 | 19.9 | 19.3 | 19.2 | 18.1 | 11.6 |

| | **Br all** | **Br···Phe** | **Br···Tyr** | **Br···Trp** | **Br···His** | **I all** | **I···Phe** | **I···Tyr** | **I···Trp** | **I···His** |
|---|---|---|---|---|---|---|---|---|---|---|
| **Count** | 445 | 167 | 102 | 121 | 55 | 75 | 49 | 14 | 3 | 9 |
| **Distance (Å)** | | | | | | | | | | |
| **mean** | 4.9 | 4.9 | 5 | 4.8 | 4.6 | 5.1 | 5.2 | 4.8 | NA | 4.9 |
| **median** | 4.9 | 4.9 | 5.1 | 4.9 | 4.6 | 5.1 | 5.2 | 4.7 | NA | 5.1 |
| **Q1** | 4.4 | 4.4 | 4.7 | 4.4 | 4.3 | 4.8 | 4.9 | 4.6 | NA | 4.6 |
| **Q3** | 5.4 | 5.4 | 5.4 | 5.3 | 5 | 5.7 | 5.8 | 5.0 | NA | 5.7 |
| **MAD** | 0.5 | 0.5 | 0.4 | 0.5 | 0.3 | 0.5 | 0.4 | 0.3 | NA | 0.5 |
| **θ1 (˚)** | | | | | | | | | | |
| **mean** | 128.5 | 133.9 | 132.0 | 122.6 | 118.7 | 131.5 | 130.8 | 146.2 | NA | 112.7 |
| **median** | 133.4 | 134.1 | 137.0 | 133.2 | 112.5 | 136.8 | 136.8 | 154.0 | NA | 110.9 |
| **Q1** | 106.1 | 111.6 | 116.7 | 102.4 | 98.8 | 111.2 | 110.8 | 150.9 | NA | 106.4 |
| **Q3** | 145.4 | 155.8 | 145.8 | 135.3 | 141.2 | 147.6 | 146.1 | 157.8 | NA | 112.1 |
| **MAD** | 18.4 | 22.0 | 14.4 | 13.5 | 15.9 | 20.8 | 11.6 | 3.7 | NA | 4.5 |
